# Substrate viscoelasticity regulates fibroblast adhesion and migration

**DOI:** 10.1101/2025.03.20.644304

**Authors:** Neha Paddillaya, Akshar Rao, Anshul Shrivastava, Imnatoshi Jamir, Kundan Sengupta, Namrata Gundiah

## Abstract

Mechanical properties of the extracellular matrix (ECM) modulate cell-substrate interactions and influence cellular behaviors such as contractility, migrations, and proliferation. Although the effects of substrate stiffness on mechanobiology have been well studied, the role of ECM viscoelasticity in fibrotic progression remains less understood. To examine how viscoelasticity affects the biophysical properties and regulates the signaling of human mammary fibroblasts, we engineered elastic (E) and viscoelastic (VE) polyacrylamide hydrogels with comparable storage moduli (∼14.52 ± 1.03 kPa) but distinctly different loss moduli. Fibroblasts cultured on E hydrogels spread extensively (2428.93 ± 864.71 μm²), developed prominent stress fibers with higher zyxin intensity, and generated higher traction stresses (2931.57 ± 1732.61 Pa). In contrast, fibroblasts on VE substrates formed smaller focal adhesion areas (54.2% reduction), exhibited lower critical adhesion strengths (51.8%), and generated 21% lower traction stresses (p < 0.001), indicating weaker adhesions. These substrates also promoted migrations and showed enhanced proliferation accompanied by reduced YAP activity, suggesting a mechanotransduction shift that may involve alternative signaling pathways. In contrast, E substrates showed YAP nuclear translocation, consistent with greater cytoskeletal tension and contractility. These findings highlight the importance of energy dissipation mechanisms in regulating fibroblast function on substrates mimicking the fibrotic milieu. Our results demonstrate that tuning the ECM viscoelasticity is a useful strategy to regulate cell behaviors in tissue engineered scaffolds, and develop better disease modeling for regenerative medicine.

## Introduction

Fibrotic tissues are characterized by excessive collagen deposition and crosslinking due to fibroblast activity, leading to a heterogeneous extracellular matrix (ECM) organization and elevated stiffness. Over time, increased tissue stiffness contributes to organ impairment and dysfunction^1,2^. A major factor in this process is fibroblast migration, which helps remodel the ECM and stiffen tissues. However, the changing ECM mechanics does not only involve increased elastic modulus but also changes in viscoelastic properties which remain less explored in the context of fibrosis. While the elastic modulus represents the immediate, reversible response of a material to stress, viscoelastic materials exhibit both solid-like elasticity and fluid-like viscous behaviors due to time-dependent responses. This viscoelasticity in the ECM, due to crosslinks and hydration, enables tissues to dissipate energy, creating a dynamic environment for cellular interactions.

The stiffness, often used as a proxy for the elastic modulus, denotes the immediate and reversible deformation response of a material; fibrotic tissues have higher stiffness. For example, breast carcinomas show elevated stiffness from 400 Pa in normal tissues to over 5 kPa in invasive ductal tissues^3^. Similarly, lung stiffness increases from 2 kPa to 17 kPa^4,5^, liver tissues from 3 kPa to 22 kPa^6^, and human atherosclerotic coronary vessels from 1.5 MPa to 3.8 MPa^7^ due to fibrotic conditions. Tsochatzis et al. showed that the optimal liver stiffness value threshold for fibrosis is 15 ± 4.1 kPa^8^. However, the stiffness does not alone capture the biomechanical milieu: tissues like the brain, liver, spinal cord, and fat show loss moduli that are 10–20% of their storage moduli across a broad range of time scales^9–11^. Studies show that patients with Type 2 diabetes mellitus and liver fibrosis have increased advanced glycation end products that alter the collagen structure leading to greater viscous dissipation without affecting the overall tissue stiffness^12^. In contrast, increasing the collagen crosslinking alone lowers the viscous contributions, resulting in myofibroblast activation^13,14^.

The viscoelastic ECM microenvironment modulates emergent cellular behaviors, such as adhesions, migration dynamics, and mechanotransduction processes, that regulate cell growth and remodeling^15–18^. Higher substrate viscoelasticity also alters the focal adhesion dynamics^17^. Human mesenchymal stem cells (hMSCs) on collagen-coated polyacrylamide gels, fabricated with similar stiffness but greater loss moduli (G’’ = 1–130 Pa at 0.005 rad/s) show increased hMSC migration, adhesion, cell spreading and differentiation into multiple lineage^19^. Enhanced filopodia formation on such substrates enhanced cell migrations^16^. Additionally, fibroblasts and cancer cells do not spread on soft, elastic alginate gels, but spread on soft, viscoelastic substrates through β1 integrin, myosin, and Rho pathways^20^. YAP nuclear translocation, a key mechanotransduction marker, is higher on elastic substrates, and correlates with stronger adhesions and proliferation, whereas cells on viscoelastic substrates favor proliferation through alternative pathways^15,21–23^.

An ideal elastic hydrogel is formed through complete covalent crosslinking, whereas crosslinked polyacrylamide (PAAm) gels, formed just beyond the gelation point, contain unbound polymer chains, leading to energy dissipation. Fabrication of gels that maintain a near constant storage modulus while exhibiting different loss moduli by varying the concentrations of the acrylamide monomer and bisacrylamide crosslinker, or by adding non-crosslinked linear acrylamide chains into the network are hence attractive to investigate mechanobiological processes^24^. We fabricated PAAm hydrogels with near similar elastic properties but significantly different viscous losses (Figure 1A). Human mammary fibroblasts (HMF3s), cultured on the elastic and viscoelastic hydrogels, exhibited distinct differences in the adhesion dynamics, tractions stresses, and migrations. YAP nuclear translocation was higher on E substrates and correlated with increased tractions compared to VE substrates. An understanding of how changes in viscoelasticity influence cellular behaviors is useful for developing targeted therapeutic strategies in fibrotic diseases, including cancers^17,25^.

**Figure 1:**
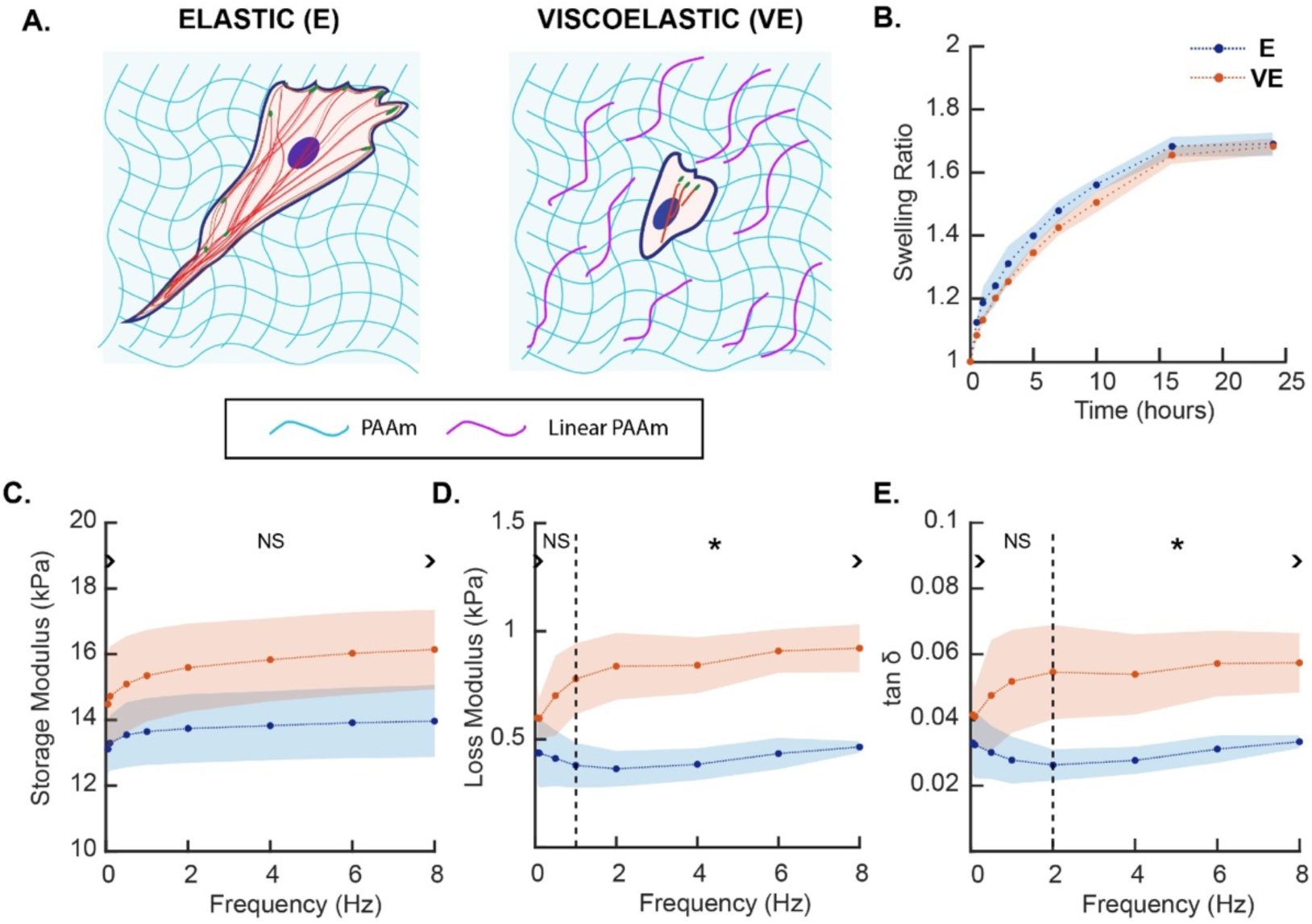
Dynamic mechanical characterization of E and VE hydrogels. (A) Schematic representation shows a top view of cells on hydrogel substrates, with entrapped linear acrylamide in VE hydrogels. (B) Time-dependent swelling of E and VE hydrogels show equilibrium properties at 15 hours. There were no differences between both groups at all time points. Variations in (C) Storage modulus, (D) Loss modulus, and (E) tan 𝛿 = 𝐸^′^/𝐸^′^ in the E and VE hydrogels are shown.

## Results and Discussion

### Fabrication and characterization of biomimetic elastic and viscoelastic hydrogels representing the fibrotic milieu

We fabricated elastic (E) and viscoelastic (VE) PAAm hydrogels by incorporating linear polymerized acrylamide into crosslinked networks (Figure 1A). This method allows us to alter the viscous contributions in hydrogels while maintaining the elastic contributions (see Materials and Methods). To assess swelling responses, we placed E and VE hydrogels in saline and periodically monitored their wet weights over 24 hours (Figure 1B). Results show marginal differences in their swelling behaviors between E (n=3) and VE (n=3) hydrogels at all measurement time points. Equilibrium swelling, defined as the point at which no further changes occurred between consecutive measurements, was reached after ∼15 hours. These results suggest that the inclusion of the viscous polyacrylamide component does not significantly alter swelling behaviors or compromise the mechanical stability of the hydrogel with time.

We conducted dynamic mechanical analysis (DMA) to assess the role of viscous inclusions on E and VE hydrogel mechanics. The storage modulus (E’), representing elastic contributions to the complex modulus, and loss modulus (E’’), reflecting viscous dissipation, were measured for the E (n=5) and VE (n=5) hydrogels over a frequency range from 0.01 to 70 Hz (Figure S1). Figures 1C and D show the mean values of these parameters over a smaller frequency range (∼8 Hz). There were no significant differences (p > 0.01) in the storage moduli between E and VE gels at loading frequencies below 18 Hz, indicating similar elastic properties under low and quasi-static conditions (Figure 1C). However, the loss modulus of VE gels was significantly higher (p < 0.01) than that of E gels at loading frequencies of 1 Hz and above (Figure 1D), demonstrating significantly higher viscous dissipation in VE hydrogels even at low frequencies. The ratio of loss to storage modulus, given by tan δ, was significantly higher (p < 0.01) for VE hydrogels at frequencies of 2 Hz and above (Figure 1E). E hydrogels had significantly low values of tan δ (p < 0.04), demonstrating low viscous contributions. We next compared the mean storage modulus, loss modulus, and tan δ values at 0.05 Hz, a frequency representative of quasi-static monotonic compression. The mean storage modulus of VE hydrogels was 10.5% higher than the E group, whereas the loss modulus was 36.9% higher than E gels at 0.05 Hz. The corresponding tan δ value was 25.9% higher for VE gels compared to E gels. These results confirm that while E and VE hydrogels have near similar storage moduli (E gels: 13.11 ± 0.76 kPa; VE gels: 14.48 ± 1.53 kPa at 0.05 Hz; p>0.1), the VE gels exhibit significantly greater viscous losses over a physiologically relevant frequency range.

The inclusion of linear polyacrylamide into VE hydrogels allowed us to modify the viscous contributions without significantly affecting their equilibrium swelling behavior. The higher loss modulus and loss tangent (tan δ) values for VE hydrogels confirm their ability to dissipate energy under dynamic loading conditions, making them particularly relevant to mimic the viscoelastic properties of fibrotic tissues.

In recent years, there has been a growing impetus in synthesizing viscoelastic hydrogels that closely resemble biological tissues and recapitulate the cellular microenvironments. Dynamic bonds and physical interactions control the viscous dissipation in hydrogels^26^. Previous studies used non-covalent crosslinking, including hydrogen bonds and hydrophobic associations, to prepare hydrogels that displayed viscoelasticity^27,28^. Dynamic covalent bonds, such as Schiff bases, borate esters, and disulfide linkages, have also been used to fabricate hydrogels with tuneable viscoelasticity due to the reversible nature of their bonds^29–31^. However, most of these methods simultaneously increase the elasticity of the hydrogel with increased viscous dissipation. Recent efforts have focussed on selectively tuning the viscous dissipation in hydrogels while maintaining nearly invariant elastic properties. Using dextran to alter the viscoelastic relaxation, Cacopardo and coworkers altered the viscosity of the hydrogel liquid phase of agarose and polyacrylamide hydrogels without changing their elastic properties^32^. Charrier and coworkers used an alternate approach and modified the solid polymer network by incorporating high molecular weight linear polyacrylamide chains into a crosslinked network to selectively tune the viscous properties^24,33^. In this method, the linear PA chains are not chemically bonded to the network but are sterically confined within its mesh, contributing to the viscous component of the hydrogel. They demonstrated the ability to generate gels with similar elasticity ranging from 0.1 to 300 kPa but variable loss modulus, by adjusting the amounts of acrylamide, bis-acrylamide, and linear PA. Such hydrogels are also suitable for cell culture and can be selectively functionalized by attaching adhesion proteins to either the crosslinked network or the linear PA chains^24,33^. Building on this approach, we prepared tuneable viscoelastic hydrogels to represent a stiffer fibrotic tissue environment. These hydrogels enable the study of viscous contributions, independent of stiffness in cell substrate interactions.

### Strength of cell adhesions on E and VE hydrogel substrates

Because viscoelastic substrates exhibit higher energy dissipation compared to purely elastic substrates, we hypothesized that higher viscous losses in VE substrates would impair cellular adhesion strength. We measured the critical adhesion strengths of human mammary fibroblasts (HMF3s) on glass (G), elastic (E), and viscoelastic (VE) hydrogels using a custom microscope-mounted fluid shear device to investigate the influence of substrate viscoelasticity on cell-substrate interactions (see Materials and Methods^34,35^). Specifically, we measured the fraction of cells adhered to the E and VE hydrogel surfaces by gradually increasing the shear stress in the device from 1.2 Pa to 8 Pa in steps of 0.2 Pa/ min. Results for the normalized cell count with increasing shear stress showed sigmoidal responses across glass (n=369), E (n=361), and VE (n=388) hydrogel substrates (Figure 2A). The critical adhesion strength (τ₅₀), defined as the shear stress required to detach 50% of cells, was significantly lower on VE hydrogels (2.15±0.08 Pa) compared to E hydrogels (4.15±0.13Pa) or glass substrates (3.81±0.16 Pa) (Figure 2B). There was however no difference in τ_50_ values measured in our study between the E and glass substrates, respectively.

**Figure 2:**
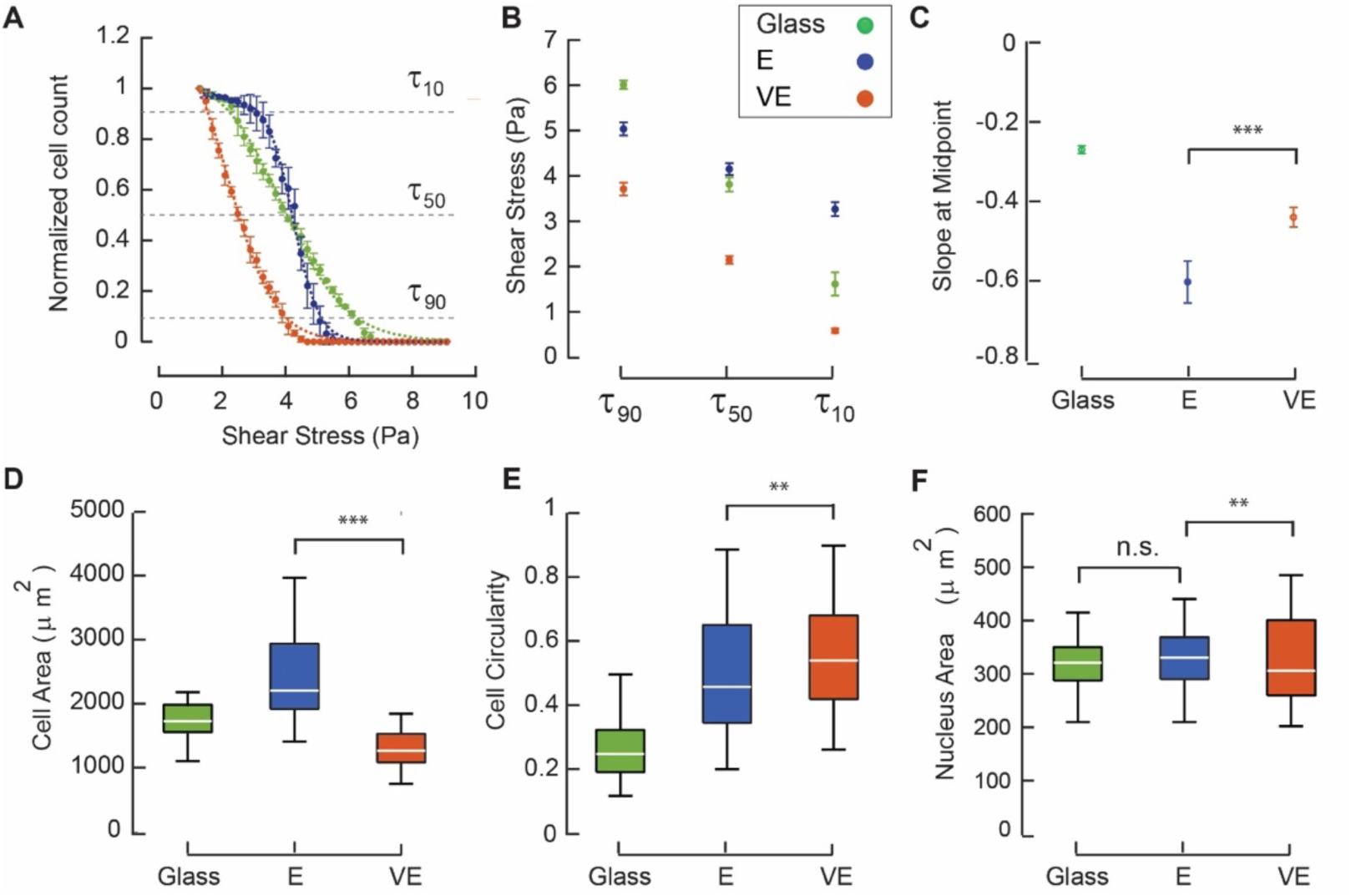
Adhesion strength and morphological differences of fibroblasts (HMF3s) on glass, E, and VE hydrogels. (A) Deadhesion curves indicate a fraction of adherent cells on substrates under increasing shear stress. (B) Critical adhesion strengths at 1_10_, 1_50,_ and 1_90_ indicating the fractions of weakly, intermediate, and strongly adherent cells are shown for all substrates. (C) Slopes of the deadhesion curves were calculated at 1_50_ and compared across the different groups. Immunofluorescence images of cells on G, E, and VE hydrogels were used to calculate the (D) Projected cell attachment area, (E) Cell circularity, and (F) Nuclear areas on E, VE hydrogels, and glass substrates in our study (p<.001, ANOVA-Bonferroni).

To further dissect adhesion dynamics on all substrates in our study, we analyzed differences in τ₁₀, the shear stress required to detach 10% of cells, which represents the contribution of weaker, transient, and flexible adhesions. Figure 2B shows significantly lower 1_10_ values on VE hydrogels (0.57±0.05 Pa) compared to E hydrogels (3.27±0.15 Pa) and glass substrates (1.62±0.26 Pa). Additionally, there was a distinct difference at τ_90,_ representing the fraction of strongly adhered cells to the substrates. E hydrogels displayed a sharp, switch-like response, indicative of a uniform density of stable integrin-mediated adhesions, in contrast to VE hydrogels and glass substrates that showed longer tail regions. To quantitatively assess adhesion heterogeneity, we compared the slopes of the detachment curves on all substrates. Cell attachments on E hydrogels displayed a steeper slope at τ_50_ (−0.60±0.05 Pa⁻¹), indicating homogenous adhesions (Figure 2C). In contrast, VE hydrogels displayed a moderately steep slope (−0.44±0.02 Pa⁻¹), reflecting a higher degree of heterogeneity on the viscoelastic substrate, whereas the glass substrates exhibited the lowest slope (−0.27±0.01 Pa⁻¹). These results demonstrate that cells on glass substrates form variable cell-substrate interactions without mechanical feedback from a deformable matrix, resulting in a distinct attachment mode compared to both E and VE hydrogel surfaces.

We analyzed differences in cell spreading on all substrates using immunofluorescence studies to corroborate results from the de-adhesion experiments (see Materials and Methods). Figure 2D shows a greater cell spread area on E hydrogels (2428.93 ± 864.71 μm², n=43) compared to VE hydrogels (1296.73 ± 311.62 μm^2^, n=40), with intermediate values observed on glass substrates (1792.61 ± 487.09 μm², n=45). This result aligns with observations that show elastic substrates support robust focal adhesion maturation and greater cytoskeletal tension^24^. In contrast, the lower spread area on VE hydrogels suggests a reduced ability to form large, stable adhesions and traction stresses. Although cell circularity did not differ significantly between VE (0.55 ± 0.16) and E hydrogels (0.50 ± 0.18; p<0.01), cells on glass were highly polarized (0.27 ± 0.11) (Figure 2E). In contrast, the nuclear areas remained unchanged across E, VE, and G substrates (Figure 2F). These findings align with previous reports on cancer cells in viscoelastic environments (0.8-5 kPa) which exhibit weak, flexible adhesions that facilitate migration and invasion^36^. The higher τ₅₀ on E hydrogels suggests that elastic substrates support stronger and mature focal adhesions, whereas VE hydrogels promote weaker and transient adhesions that are more susceptible to detachment. The enhanced adhesion strength and spreading on elastic hydrogels are consistent with the reinforced focal adhesions and elevated cytoskeletal tension typical of cancer-associated fibroblasts (CAFs) and tumor cells in stiffer microenvironments caused by increased ECM deposition seen in fibrotic conditions^25^. These differences between adhesions on E and VE hydrogels may be useful in understanding how cancer cells transition from stable adhesions in less viscous microenvironments to flexible, transient adhesions that facilitate invasion in microenvironments with greater viscosities.

#### Traction stresses on elastic and viscoelastic hydrogel surfaces

Cells spread extensively on stiffer matrices and generate higher traction stresses through actin stress fibers and focal adhesion complexes^24^. We hypothesized that the elastic substrates facilitate the formation of stronger stress fibers, leading to higher tractional stresses compared to VE substrates. We performed traction force microscopy (TFM) using a regularized FTTC (Reg-FTTTC) approach on HMF3s fibroblasts seeded on E and VE hydrogels (see Materials and Methods)^37^. Brightfield images of cells on E and VE hydrogels (Figure 3A) show distinct morphological differences. Representative images of the displacement maps and corresponding maximum tractions illustrate the influence of substrate viscoelasticity on cellular contractility. Figure 3B shows that fibroblasts on E substrates generated significantly higher traction stresses (2931.57 ± 1732.61 Pa, n=19) compared to those on VE substrates (617.27 ± 177.97 Pa; n=20 p < 0.05). The total strain energy (Figure 3C) showed a similar trend: cells on E substrates generated significantly greater strain energy (0.19 ± 0.12 pJ) compared to those on VE substrates (0.01 ± 0.01 pJ; p<0.05). The net contractile moment (Figure 3D) was also significantly higher on E substrates (5.97 ± 4.26 Pa·μm) compared to VE substrates (0.95 ± 0.57 Pa·μm; p<0.001). The magnitude of tractions depend on the cell type: for example, 3T3 fibroblasts exert tractions ranging from 0.25 to 0.5 kPa on PAAm substrates with stiffness between 2.8 and 30 Pa^38^. These results are consistent with those reported in other studies demonstrating that viscoelastic environments promote dynamic cytoskeletal remodeling, characterized by increased actin turnover, reduced stress fiber stability, and reduced focal adhesion size^39,40^.

**Figure 3:**
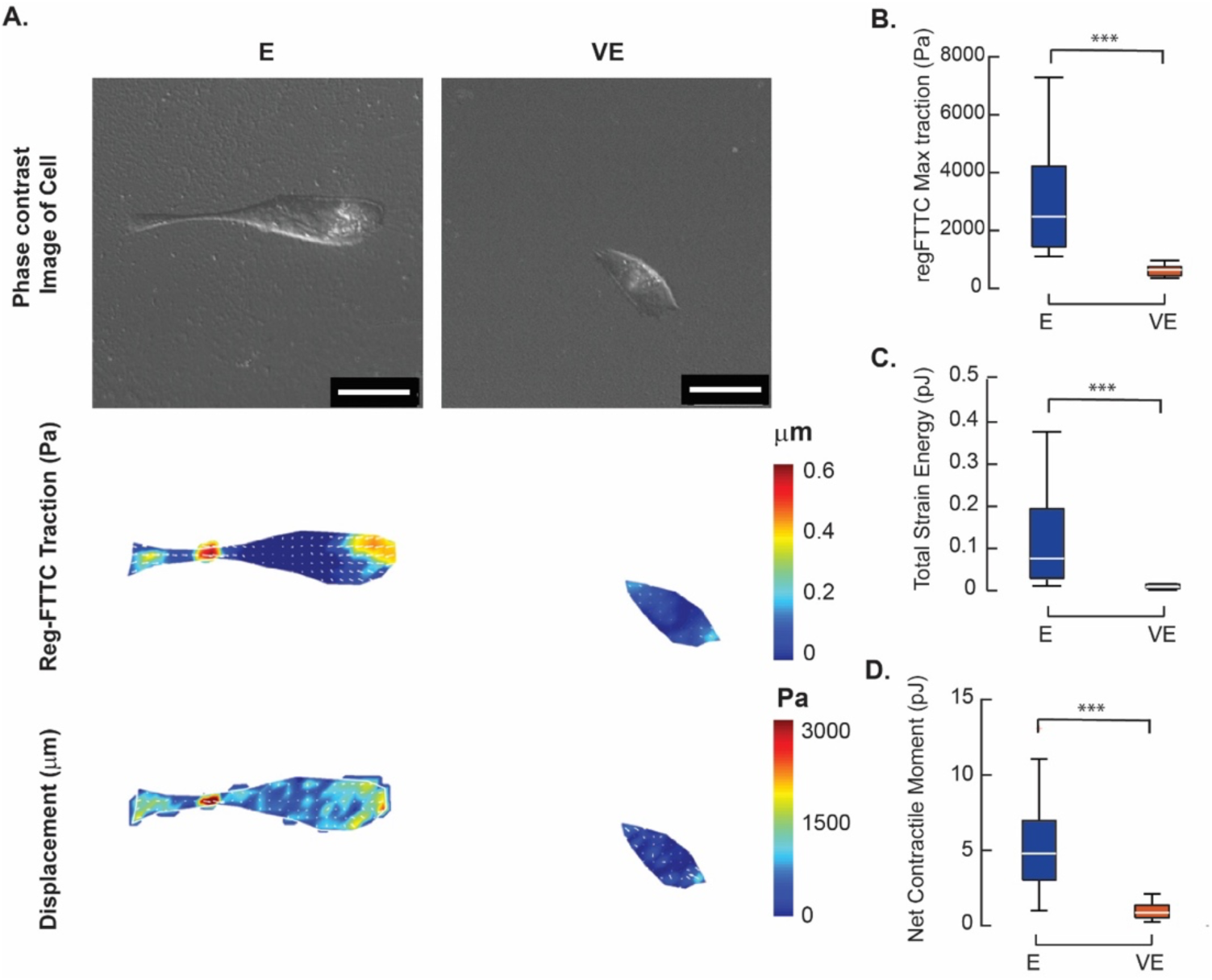
Traction Force Microscopy (TFM) of Fibroblasts on Elastic (E) and Viscoelastic (VE) Hydrogels. (A) Phase contrast images of HMF3s fibroblasts are shown on E and VE hydrogels. Corresponding displacements and Reg-FTTC traction stress maps are shown. (B) Maximum Reg-FTTC traction stress (Pa), (C) Total strain energy (pJ), and (D) Net contractile moment generated by cells on E hydrogels compared to VE hydrogel substrates are shown. (n=24,27) (p<.001, ANOVA-Bonferroni).

Cells on stiffer elastic substrates form stronger focal adhesions and generate higher actomyosin contractile forces^41^. Stiff substrates promote the assembly of stress fibers and reinforce focal adhesion complexes, including vinculin, talin, and zyxin, enabling cells to exert elevated contractile stresses on their environment. Increased traction stress and strain energy on stiffer, elastic substrates reflect the role of substrate elasticity in promoting stable adhesions and robust mechanotransduction. In contrast, viscoelastic substrates promote dynamic and transient interactions, potentially mimicking the remodeling environments in tissues.

We next tested the presence of actin in cells on E and VE substrates. Stress fibers, composed of F-actin bundles, crosslinked by α-actinin, and powered by non-muscle myosin II, play a key role in regulating cell shape, contractility, and adhesion stability^17,41^. Figure 4A shows representative immunofluorescence images of HMF3s fibroblasts stained for actin (red), zyxin (green), and DAPI (blue). The average actin intensity per unit cell was significantly higher on E hydrogels (23.18 ± 4.81 AU; n=11), followed by VE hydrogels (15.33 ± 2.83 AU; n=13); the lowest intensity was observed on glass substrates (12.21 ± 2.58 AU; n=12; p<0.01) (Figure 4B).

**Figure 4:**
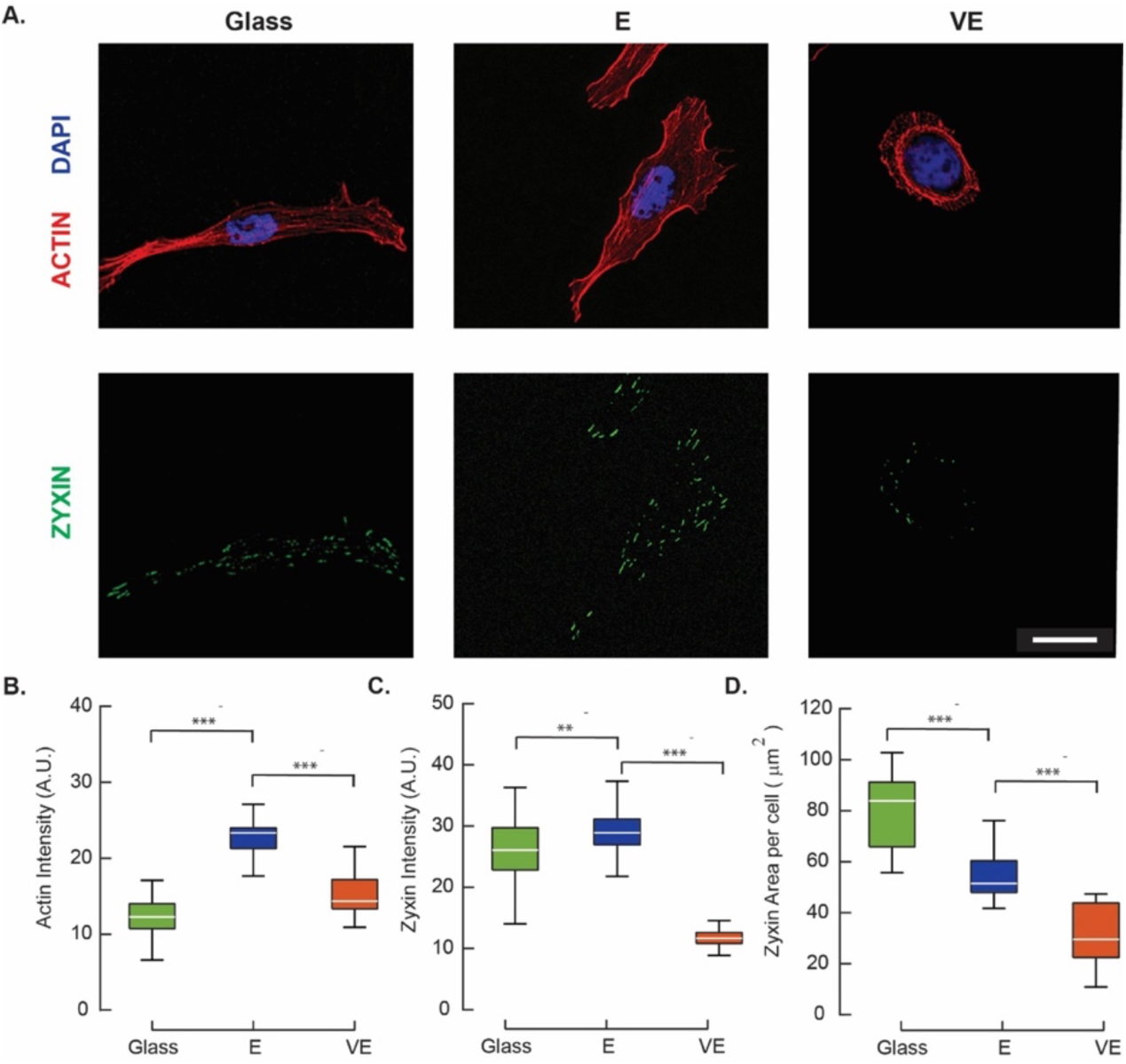
Stress fiber and focal adhesion formations on substrates. (A) Immunofluorescence images show actin stress fibers (red), zyxin (green), and nuclei (blue), on Glass, E, and VE substrates. (B) Mean actin intensity per cell, (C) Zyxin intensity per cell, and (E) Zyxin area per cell are shown for fibroblasts on substrates. (p<0.001)

We used these images to analyze the presence of zyxin — a key focal adhesion protein that facilitates mature adhesion stabilization — to link the reduced cellular traction stresses (Figure 3B) and lower adhesion strengths (Figure 2A, 2B) on VE substrates with impaired focal adhesion maturation^42,43^. Zyxin intensity per cell (Figure 4D) was highest on E hydrogels (30.03 ± 5.98 AU, n=13), followed by glass substrates (25.92 ± 5.47 AU, n=15), and lowest on VE hydrogels (11.78 ± 2.18 AU, n=14). However, the average zyxin area per cell (Figure 4E) was significantly higher on glass substrates (80.37 ± 14.22 μm²) compared to E (55.98 ± 17.64 μm²) and VE hydrogels (30.34 ± 11.56 μm²; p < 0.001). These results indicate that stiff substrates promote robust focal adhesion formation and enhance the mechanical reinforcement between the ECM and the cytoskeleton.

Riveline and coworkers (2001) showed that focal adhesion size and stability increased on stiffer substrates due to enhanced mechanical linkages between the ECM and the cytoskeleton in NIH3T3 and SV-80 human fibroblasts^39^. Similarly, hepatic stellate cells on viscoelastic (G” = 500 Pa) gels exhibit reduced α-SMA expression, smaller focal adhesions, and lower actomyosin forces as compared to 5 kPa elastic (G” = 0 Pa)^33^ hence these cells reverted to a non-differentiated fibroblast phenotype on viscoelastic substrates in contrast to the myofibroblast phenotype on elastic substrates. Dermal fibroblasts on slow-relaxing viscoelastic films also exhibit longer stress fibers and increased inflammatory markers, indicating a transition to a myofibroblast-like phenotype^44^. However, faster-relaxing substrates inhibit this transition, highlighting the importance of relaxation rates that regulate cell behaviors^44^. In contrast, Janmey and coworkers observed that the spreading behaviors of cells on viscoelastic polyacrylamide substrates varied based on the cell type: hepatocellular carcinoma cells (Huh7) displayed higher spreading, whereas normal hepatocytes, including primary human hepatocytes (PHH), showed reduced spreading on VE substrates (G’= 5kPa and G’’= 600Pa) compared to more elastic substrates (G’= 5kPa and G’’= 0Pa) which depend on adhesion dynamics and cytoskeletal regulation^45^. Results from our study also demonstrate that smaller and more dynamic adhesions of HMF3s cells on VE substrates facilitate transient interactions with the ECM, allowing cells to detach and reattach more easily. Viscoelastic substrates hence enable adaptable cell-matrix interactions which promote rapid cytoskeletal remodeling that is essential in migration.

### High motility and directed migrations characterize fibroblast responses on VE hydrogels

Altered focal adhesions and contractility measured in our study on fibrotic E and VE substrates in our study highlight the importance of energy dissipation mechanisms in modulating adhesion dynamics and actin polymerization. Because VE hydrogels facilitate transient and dynamic adhesions, we hypothesized that cells on these substrates would migrate significantly faster compared to those on E hydrogels. We tracked the migration of HMF3s fibroblasts over 12 hours on E hydrogel (n=103), VE hydrogel (n=101), and glass (n=103) substrates, respectively. Trajectories of HMF3s fibroblasts on VE hydrogel showed higher displacements compared to E hydrogel and glass substrates (Figure 5A). Figure 5B shows the mean square displacement (MSD) which measures the average distance traveled by cells from their original position over time (see Materials and Methods). We used these results to calculate the exponent (α) to distinguish the different migration modes of cells on the various substrates in our study. A purely diffusive behavior corresponds to α = 1, whereas higher values of α indicate directed fibroblast migrations on substrates. α values were 1.12 on glass substrates, which increased to 1.24 on E hydrogels, and to 1.32 on VE substrates, suggesting persistent and super-diffusive migrations. The directionality ratio, a measure of the persistence of migration, also revealed clear differences between the various substrates in our study (Figure 5D). Fibroblasts on VE and E hydrogels showed increased persistence but transitioned to a Brownian-like migration after ∼5 hours. Migration speeds were also significantly elevated on VE hydrogels compared to glass and E hydrogels (Figure 5C). These data support the reduced expression of zyxin (Figure 4D), indicating that viscous substrates promote smaller focal adhesions and hence dynamic interactions with substrates, thereby facilitating higher cellular motility. Chester et al. (2018) investigated poly (N-isopropyl acrylamide)-co-(acrylic acid) microgel films (tan δ = 0.8–1.8), where increased tan δ of the substrates showed an increase in amoeboid like migration in human dermal fibroblasts^44^. These findings demonstrate that fibroblasts exhibit increased migration persistence and super-diffusive behavior on viscoelastic (VE) substrates compared to elastic (E) hydrogels and glass, with a transition to Brownian-like migration after prolonged durations which suggests that substrate viscosity plays a crucial role in facilitating dynamic cell-substrate interactions, promoting higher cellular motility.

**Figure 5:**
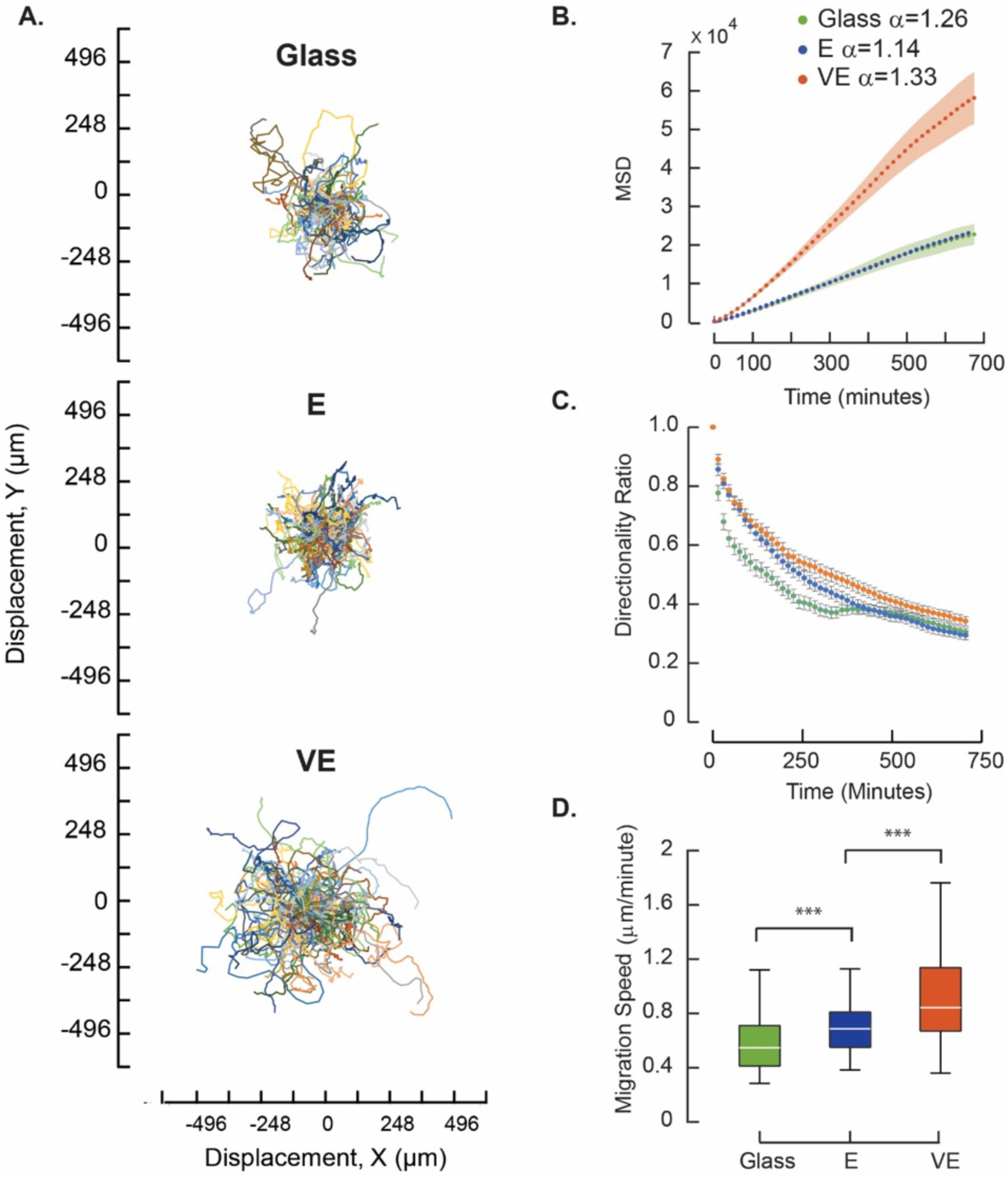
Cell migrations are different on E hydrogel, VE hydrogel, and glass substrates. (A) Fibroblast displacement tracks are shown over 12 hours. (B) MSD variations, (C) Directionality ratios, and (D) Migration speeds (p<0.001) with time show clear differences on the various substrates. Statistical analysis was performed using the Kruskal-Wallis test with Dunn’s post hoc corrections.

### YAP activation and proliferations differ on E and VE substrates

We quantified the presence of Yes-associated protein (YAP), a key transcriptional co-activator in mechanotransduction that translates mechanical cues into gene expression changes^46^. Immunofluorescence images of YAP localization (Figure 6A) reveal distinct differences between E and VE hydrogels that have comparable stiffness. Figure 6B shows a plot of the nuclear-to-cytoplasmic YAP intensity ratios that demonstrates a significantly higher value on E hydrogels (5.52 ± 0.70, n=47) compared to glass substrates (3.71 ± 0.57; n=45, p<0.001); the lowest ratio was observed on VE hydrogels (1.58 ± 0.34; n=49, p<0.001). The increased nuclear YAP on hydrogel substrates aligns with mechanotransduction, where cytoskeletal tension is transmitted to the nucleus *via* actin filaments and focal adhesion complexes^47^. Mechanical forces regulate YAP activity that modulates cell growth and division^46,48^. Actomyosin contractility compresses the nucleus, opening nuclear pores to facilitate YAP/TAZ nuclear accumulation, and amplifies mechanical stimuli. However, the nuclear localization YAP was significantly reduced on VE hydrogels, suggesting the existence of an alternative mechanotransduction pathway.

**Figure 6:**
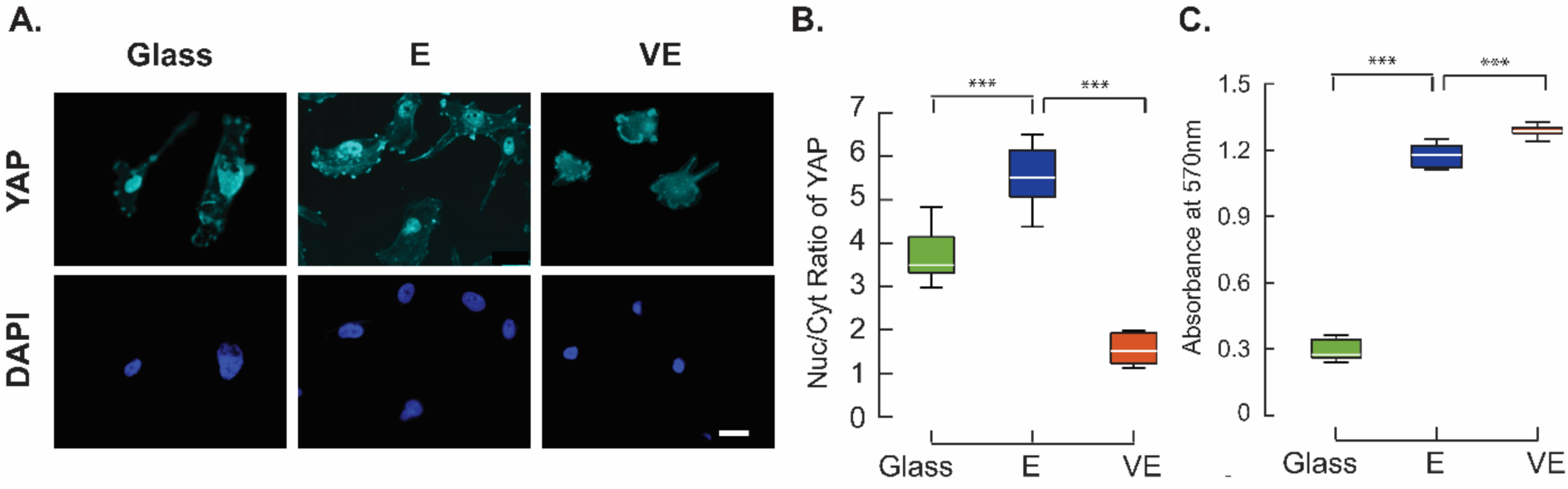
YAP translocation and proliferation in fibroblasts on substrates. (A) Representative immunofluorescence images of YAP localization are shown on glass, E, and VE hydrogels. (B) The nuclear-to-cytoplasmic intensity ratio of YAP shows clear differences on various substrates. (C) Results from the MTT assay demonstrate significantly increased proliferations on hydrogel surfaces compared to glass.

We hypothesized that these changes in YAP expressions correlated with differences in cell proliferations on the hydrogel substrates. Although YAP is known to regulate cell proliferation^46,48,49^, few studies have measured these changes on elastic and viscoelastic substrates that mimic the fibrotic condition. We quantified the proliferation of fibroblasts on E, VE hydrogels, and glass substrates using an MTT assay to measure the metabolic activity as an indicator of cell viability and growth. Our experiments show that fibroblast proliferation was significantly higher on VE hydrogels compared to E substrates (p < 0.01; Figure 6C).

Bauer et al. (2017) employed alginate RGD-modified hydrogels (E = 2.8–49.5 kPa, t₁/₂ = 79–519 s), where softer and more rapidly relaxing substrates enhanced adhesion and proliferation of C2C12 cells, but stiffer substrates increased differentiation^50^. Chaudhuri et al. (2019) demonstrated that alginate hydrogels (∼3 and ∼16 kPa) with varying stress relaxation rates promote cell proliferation in slow-relaxing gels through YAP-independent mechanisms by increasing cytoplasmic p27^Kip1^, potentially involving the PI3K-AKT pathway or integrin-independent signaling^51^. Our findings show similar cellular responses on stiffer but viscous gels, suggesting a mechanotransduction shift on VE substrates, where cells experiencing lower mechanical forces rely on alternative pathways such as PI3K-AKT rather than force-dependent YAP activation for proliferation^52^. YAP activation drives proliferation in cells that rely on biochemical cues to regulate proliferation^53,54^, or through matrix viscoelasticity, promoting liver cancer progression^36^. The cellular responses and YAP activity may also differ based on the dimensionality and composition of the matrix microenvironment^55^. The increased ability of VE substrates to dissipate mechanical energy reduces the mechanical signals required for YAP activation, shifting cellular responses toward biochemical regulation of growth and cellular adaptation^49^. These effects are hence relevant during wound healing, and in fibrosis and cancer progression. These results could have exciting therapeutic implications in targeting the YAP/TAZ-mediated mechanotransduction in treating fibrosis and mechanosensitive diseases.

### Conclusions

An understanding of how ECM mechanics regulate cellular behavior is crucial for tissue engineering and therapeutic strategies in managing fibrotic conditions. During the initial stages of fibrosis, viscoelastic effects influence cellular behavior by modulating fibroblast migration, adhesion, and activation. Increased substrate viscosity enhances fibroblast motility and promotes persistent migration, facilitating tissue remodeling. These biomechanical cell-substrate interaction cues to substrate relaxation may accelerate extracellular matrix (ECM) deposition and fibrosis progression. We demonstrate that biomimetic substrates for fibrotic tissue conditions, which are comparable in storage moduli but have significantly different loss moduli, are accompanied by markedly distinct cell-matrix interactions. The VE hydrogels promoted weaker, more dynamic adhesions, with higher and directed migrations and increased proliferation. In contrast, E hydrogels favored the formation of robust focal adhesions, and showed enhanced traction stresses and pronounced YAP nuclear translocation. These findings highlight the critical role of substrate viscoelasticity in regulating fibroblast behavior and hence fibrosis progression. Tuning of the tissue stiffness and viscoelasticity may hence be desired to accurately replicate physiological and pathological tissue mechanics, ultimately enabling more effective strategies for tissue repair, disease modeling, and high-throughput drug screening.

## Methods

### Fabrication of hydrogels to mimic the fibrotic tissue milieu

We engineered polyacrylamide hydrogels by crosslinking acrylamide with bisacrylamide in addition to using ammonium persulfate (APS) and tetramethylethylenediamine (TEMED)^56^. We combined a 40% (w/v) acrylamide stock solution with a 2% (w/v) bisacrylamide stock solution in deionized water to fabricate E hydrogels according to the concentrations specified in Table 1. We also prepared a linear polyacrylamide solution by crosslinking 1.25 mL of 40% (w/v) acrylamide with 0.025 mL of 10% (w/v) APS and 0.005 mL of TEMED in 8.72 mL of deionized water (Table 1)^24^. This mixture was polymerized for 1 hour at 37 °C to obtain a highly viscous linear polyacrylamide (Linear PAA). Viscoelastic VE hydrogels were fabricated by combining the prepared Linear PAA with acrylamide and bisacrylamide stock solutions (Table 1). Both E and VE mixtures were poured into cylindrical molds (16 mm in diameter, 10 mm in height), followed by the addition of APS (10% w/v) and TEMED (Sigma Aldrich, T9281). Each mixture was stirred using a pipette, degassed and the mixture polymerized at room temperature for 30 minutes. The hydrogels were gently removed from the molds and stored in deionized water at 4°C overnight for use in cell-based experiments.

**Table 1:** Concentrations of acrylamide, APS, and TEMED to prepare 10 ml solution for the fabrication of E and VE PAAm hydrogels.

|  | Acrylamide<br>(40% w/v) | Bisacrylamide<br>(2% w/v) | Linear<br>PA | DI<br>Water | APS<br>(10% w/v) | TEMED |
| --- | --- | --- | --- | --- | --- | --- |
| **E** | 2 ml<br>(20 %) | 0.50 ml<br>(5.0 %) | - | 7.40 ml<br>(74.0 %) | 0.075 ml<br>(0.75 %) | 0.025 ml<br>(0.25 %) |
| **VE** | 2 ml<br>(20 %) | 0.75 ml<br>(7.5 %) | 1.5 ml<br>(15 %) | 5.65 ml<br>(56.5 %) | 0.075 ml<br>(0.75 %) | 0.025 ml<br>(0.25 %) |

### Gel swelling experiments

E and VE hydrogels were weighed and next immersed in 5 mL of 1X PBS at room temperature to evaluate the swelling behaviors^34^. We weighed the wet weights with a weighing balance. Swelling ratios were calculated as:

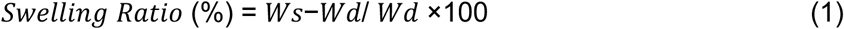

W_s_ is the weight of the swollen sample at each time point. W_d_ represents the dry weight of the sample before swelling.

### Characterization of the viscoelastic properties of hydrogels using a DMA

We used a Bose ElectroForce® 3200 dynamic mechanical analysis (DMA) system to quantify the viscoelastic properties of the engineered E and VE hydrogels. Each specimen was preloaded with 10 g force, and subjected to a sinusoidal displacement of 0% to 10% strain across frequencies ranging from 0.05 to 70 Hz. A 25 N load cell (Honeywell Sensotec Inc., Columbus, OH, USA) recorded the corresponding changes in the forces using a data acquisition rate of 30 times the loading rate. These data were used to calculate the storage and loss moduli for each loading block using a custom MATLAB code (MATLAB version: 9.13.0 (R2022b), Natick, Massachusetts: The MathWorks Inc.; 2022)^34^. Specifically, We applied Fourier transforms to the stress and strain data for each frequency block, and calculated the resulting phase and amplitude values which were used to obtain the storage (E’) and loss (E”) moduli for each specimen (n = 3 for each group). We also determined the loss tangent (tan δ = E”/E’) using these data^34^.

### Cell culture and sample preparation

Human mammary fibroblast cells (HMF3s) were cultured in DMEM (Invitrogen) supplemented with 10% FCS (Invitrogen) and 1% penicillin/streptomycin (Sigma-Aldrich) at 37°C with 5% CO₂. Cells were passaged every 2–3 days and harvested using 0.25% trypsin-EDTA (Invitrogen). To prepare cell-seeded hydrogel substrates, we treated clean 22 mm glass coverslips with 3-aminopropyl triethoxysilane (APTES) and glutaraldehyde. Acrylamide– bisacrylamide solution was next polymerized between the treated coverslip and an untreated coverslip to help attach the hydrogel to the glass. The untreated coverslip was removed after polymerization to expose the hydrogel surface. A 22 mm clean coverslip, plasma-treated for 3 minutes and coated with 50 µg/mL rat tail collagen I (Gibco), served as a control glass substrate in this study. 100 mg/mL stock of Pierce™ Sulfo-SANPAH (No-Weigh™ Format, Thermo Fisher) was diluted 1:200 in 50 mM HEPES buffer, pipetted onto the gels (200 µL), exposed to UV light for 10 minutes, and then washed to prepare E and VE hydrogel substrates. This process was repeated prior to coating with 75 µL of 50 µg/mL rat tail collagen I, which was followed by incubation at 37 °C for 45 minutes. Gels were then rinsed with PBS, seeded with HMF3s (5000 cells/mL), and allowed to attach for 6 hours before starting the experiments.

### Deadhesion studies

Nuclei of cells cultured on either glass substrates, E or VE hydrogel substrates were stained with Hoechst (Invitrogen) for 3 minutes at 37 °C in a humidified incubator (5% CO₂). Specimens were washed twice with PBS to remove excess stain, and replaced with fresh before mechanical testing. A custom microscope-mounted fluid shear device was used to measure cell de-adhesion strengths based on protocols developed earlier^57^. The prepared petridish with cells was placed on the device mounted to an inverted microscope (10×; Leica DMI6000, Germany), and the cone was set 20 µm above the sample surface using a z-translation stage. The device works on the principle of generating a Couette flow which was used to create controlled shear stress on the cells plated on glass, E and VE hydrogel surfaces. The fluid shear stress was ramped from 1.2 Pa, at 0.2 Pa each minute using a custom Arduino script. Fluorescence images were captured at each shear stress increment using a CCD camera (DFC 365X). The images were next analyzed in FIJI^58^ to count the cells at each increment of shear. These data were used to measure the fraction of cells on the substrate with increasing shear stress. The critical de-adhesion strength was defined as the stress at which 50% of cells remained. τ_10_ and τ_90_ were also calculated and compared between the various substrates in our study.

### Traction force microscopy (TFM)

A Regularized Fourier Transform Traction Cytometry (reg-FTTC) method ^37^ was used to calculate traction stresses exerted by cells on the E and VE hydrogels. Specimens were prepared on two sets of 22 mm glass coverslips. The first set was surface-treated with 3-aminopropyl triethoxysilane (APTES) and glutaraldehyde, and the second set was treated with 1% BSA. Fluorescent polystyrene beads solution (200 nm, Thermo, 16:1000) was spin-coated onto the BSA-treated coverslips. An acrylamide-bisacrylamide solution was next polymerized between the two coverslips, and the gels functionalized using 1:200 Pierce™ Sulfo-SANPAH (100 mg/mL stock, Thermo) at 302nm UV, and coated with 50 µg/mL rat tail collagen I (Gibco). HMF3s fibroblasts (2000 cells/mL) were seeded on these gel substrates, and allowed to attach for 6 hours before experiments. Brightfield images of cells and fluorescence images of homogenously coated beads on hydrogel surfaces were obtained using a live-cell microscope (40× oil immersion; Leica DMI6000, Germany) equipped with a PECON stage incubator. Cells were next removed from hydrogel substrates using SDS, and a second fluorescent bead image was acquired to capture the referential configuration of the gel. The deformed and referential epifluorescence bead images were used to compute the displacement fields and the regularized FTTC tractions were calculated using a custom MATLAB Reg-FTTC program^37^.

### Cell migration experiments

HMF3 cells (5000 cells/mL) were seeded on G, E, and VE hydrogel substrates for migration studies. Time-lapse experiments were performed 6 hours after cell sedding using an inverted live-cell epifluorescence microscope (Leica DMI6000B, Germany) at 10× phase contrast. Images of single cells during migration were captured every 15 minutes over 12 hours, with ∼10 positions imaged per condition using a motorized stage. These images were imported into Fiji^58^, converted to 8-bit, and drift-corrected using a Template Matching plugin (bilinear interpolation), and the cells were tracked using a Manual Tracking Plugin (MTrackJ) plus centering correction. All tracks were aligned to a common origin in the final plot.

Trajectories from the glass substrate, E and VE hydrogels were used to calculate the Mean Squared Displacement (MSD) and directionality ratio as described earlier^59^. MSD follows a power-law dependence with time for active Brownian particles, and is given by:

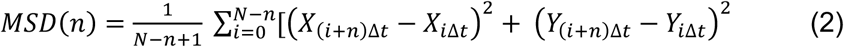

Where MSD(𝑛) is the mean squared displacement at step size 𝑛, with Δ𝑡 set by the frame rate and 𝑁 as the total displacements per trajectory. The slope (𝛼) of log-log MSD plots shows directional persistence. Population averages are calculated equally across all cells, without prioritizing longer trajectories, focusing on short-time intervals to determine α-values. Values of 0<𝛼<1 indicate subdiffusion, while 𝛼>1 is superdiffusive^60^. Cell speeds were next obtained and the directionality ratio was calculated as:

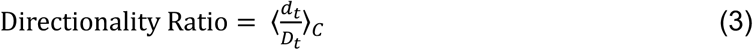

d_t_ is the straight-line distance from the start point to the cell’s position at time 𝑡. 𝐷_𝑡_ is the actual trajectory length over the same interval. The DR ratio was used to evaluate the straightness of the cellular trajectories.

### Immunofluorescence studies

Cells were fixed in 4% paraformaldehyde pH 7.2, permeabilized (0.1% Triton X-100, 3 min), blocked (5% BSA + 5% FBS, 1 h at RT), and incubated for 1 h with anti-zyxin and anti-YAP (1:400, Sigma). Next, Alexa Fluor 488 secondary antibodies (1:800) and Rhodamine phalloidin (1:200) were applied for 1 h at RT, with PBS washes after each step. DAPI (1:500) labeled nuclei for 2 min, and specimens were mounted (Goldfade, Invitrogen) before imaging on an Andor Dragonfly Spinning Disc microscope (IISc Bioimaging Facility). Images were analyzed in Fiji and were used to compute the cell area, circularity, and aspect ratio.

### MTT proliferation assay

MTT assay was done in a 24-well plate. Each well contained 5000 cells/mL on glass, E or VE hydrogel substrates (total 500 µL). Empty wells served as MTT blanks. The wells were incubated for 24 hours at 37 °C (5% CO₂), and 50 µL MTT (5 mg/mL) was added into each well. The media was removed after 4 hours followed by 3 PBS washes, and 300 µL DMSO was added into each well to dissolve formazan crystals. Plates were placed on a shaker (10 min) for complete dissolution, and the final absorbances were read at 570 nm to quantify the cell proliferations.

### Statistical analysis

Data were analyzed in GraphPad Prism 8.4.2 and presented as mean ± SD or SEM. Comparisons between two groups were performed using two-tailed unpaired Student’s t-tests, whereas one-way ANOVA with Bonferroni’s post hoc tests covered multiple groups. Normality was checked with the Shapiro-Wilk test (95% CI). For non-normal data in migration analysis, Kruskal-Wallis with Dunn’s post hoc tests were applied. All statistical details are given in figure captions; significance * p<0.5, ** p<.01 and *** p<.001.

## Author contributions

NP, AR: Methodology, Software, Validation, Formal Analysis, Investigation, Data Visualization. AS, IJ: Material Characterization. NP: Writing – Original Draft, Editing, KS: Conceptualization, Writing – Reviewing and Editing, Supervision, and Funding Acquisition. NG: Conceptualization, Methodology, Software, Formal Analysis, Resources, Data Curation, Writing – Original draft, Reviewing and Editing, Data Visualization, Supervision, Project Administration and Funding Acquisition.

## Acknowledgments

We are thankful to the Bioimaging Facility, Indian Institute of Science (IISc) for the use of the Andor Dragonfly 502 spinning disk confocal microscope. NG and KS are grateful to MoE-STARS (2/2023-0603) for project support.

